# The role of alternative polyadenylation in Epithelial-mesenchymal transition of non-small cell lung cancer

**DOI:** 10.1101/2023.11.02.565398

**Authors:** Sijia Wu, Xinyu Qin, Liyu Huang

## Abstract

The metastatic non-small cell lung cancer (NSCLC) is one of the cancers with high incidence, poor survival, and limited treatment. Epithelial-mesenchymal transition (EMT) is the first step by which an early tumor converts to an invasive one. Studying the underlying mechanisms of EMT can help the understanding of cancer metastasis and improve the treatment. In this study, 1,013 NSCLC patients and 123 NSCLC cell lines are deeply analyzed for the potential roles of alternative polyadenylation (APA) in the EMT process. A trend of shorter 3’-UTRs is discovered in the mesenchymal samples. The identification of EMT-related APA events highlights the proximal poly(A) selection of *CARM1*. It is a pathological biomarker of mesenchymal tumor and cancer metastasis through losing miRNA binding to upregulate the EMT inducer of *CARM1* and releasing miRNAs to downregulate the EMT inhibitor of *RBM47*. The crucial role of this APA event in EMT also guides its effect on drug responses. The patients with shorter 3’-UTR of *CARM1* are more benefit from chemotherapy drugs, especially cisplatin. A stratification of NSCLC patients based on this APA event is useful for chemotherapy design in future clinics.

## Introduction

Lung cancer remains the leading cause of cancer death [1]. Its main subtype is non-small cell lung cancer (NSCLC), constituting 85% of lung cancer patients [2]. About 30–40% of all people who receive an NSCLC diagnosis have metastatic cancer [3]. Epithelial-mesenchymal transition (EMT) provides stationary epithelial cells with the ability to migrate [4]. It allows solid tumors to become more malignant, increasing their invasiveness and metastatic activity [5-6]. The further exploration of EMT mechanisms of NSCLC can help understand cancer metastasis and improve NSCLC treatment.

Previous studies reveal a series of molecules involving the EMT process [7]. For example, *TWIST* is a transcription factor promoting EMT [8]. The dysregulation of proteins, such as N-cadherin and E-cadherin, is one hallmark of EMT [9]. The *miR-200* family participates in the TGF-β-induced EMT and affects drug resistance [10]. The activation of NF-κB signaling induces EMT to enhance the metastasis of tumor cells [11]. Due to the refractory metastatic NSCLC, more understanding of the molecular mechanisms of EMT is required [12].

Alternative polyadenylation (APA) can be a promising target for inhibiting EMT and metastasis [13]. The proximal selection of poly(A) site leads to the depletion of one APA subunit, stimulating EMT and increasing migration [13]. One APA writer, *CSTF2*, induces 3’-UTR shortening and upregulates a stimulator of EMT, *FGF2* [14]. The shorter transcript of *ZEB1*, one EMT-related transcription factor, is associated with chemotherapy resistance [15]. The potential of APA in EMT and its limited research in lung cancer guide us to this study.

To do this, we first define epithelial or mesenchymal phenotype for each NSCLC sample. After a comparison of APA events between epithelial and mesenchymal groups, we observe the preferred selection of proximal poly(A) site after the transition from epithelial to mesenchymal status. Based on all the differential APA candidates, 14 APA biomarkers highly associated with EMT are identified. They show an effective prognosis ability of EMT status. Of them, one APA biomarker is studied deeply for its potential roles and mechanisms to affect EMT and drug sensitivity.

## Results

### 1. Shortened 3’-UTRs in mesenchymal tumors of NSCLC

Our pipeline for the evaluation of EMT status identifies 706 epithelial samples and 104 mesenchymal lung cancer patients from The Cancer Genome Atlas (TCGA). Similarly, there are 53 cell lines showing epithelial characteristics and 49 cell lines displaying mesenchymal features from Cancer Cell Line Encyclopedia (CCLE). The two groups are clearly separated using the first two principal components of the expressions of EMT genes (Figure 1A). The mesenchymal samples get higher scores for the three EMT-related pathways compared to the epithelial ones (Figure 1B-C). The results are consistent in both TCGA and CCLE datasets, revealing the reliability of our pipeline for the classification of epithelial and mesenchymal phenotypes.

**Figure 1.**
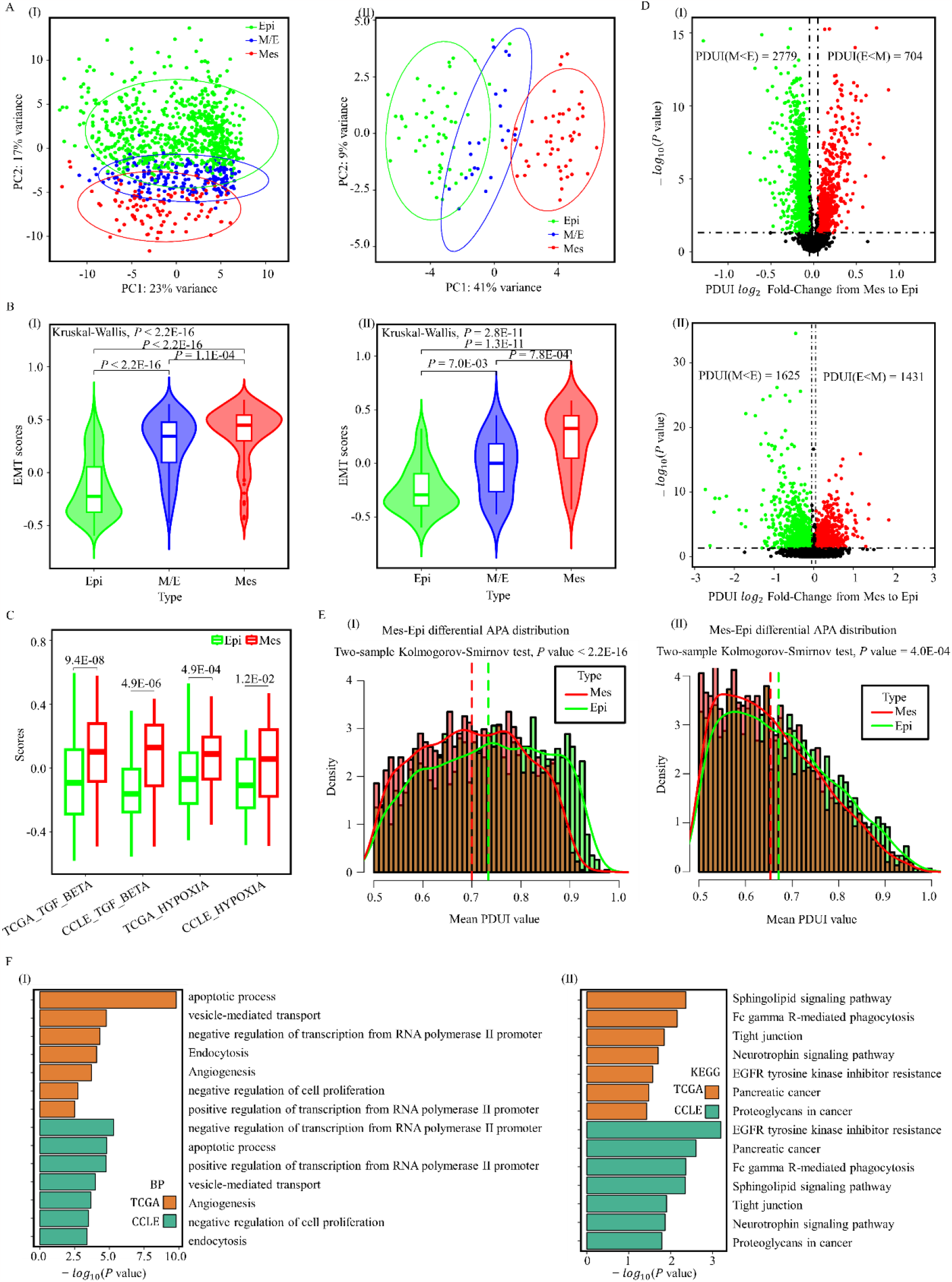
The overview of alternative polyadenylation in mesenchymal lung cancer samples. (A) PCA plots showing the definition of epithelial and mesenchymal patients from TCGA (left) and cell lines from CCLE (right). (B) The comparison of EMT gene set score between the defined epithelial and mesenchymal patients (left) and cell lines (right). (C) The comparison of two EMT-related gene set scores between the defined epithelial and mesenchymal samples. (D) The comparison of APA events between the defined epithelial and mesenchymal patients (top) and cell lines (bottom). (E) The PDUI distributions of average differential APA events for epithelial and mesenchymal patients (left) and cell lines (right). (F) The enriched biological processes (left) and KEGG pathways (right) of genes with differential APA events for TCGA patients and CCLE cell lines. PCA, Principal component analysis; Epi, epithelial samples; Mes, mesenchymal samples; M/E, the unclassified samples; APA, alternative polyadenylation; PDUI, distal poly(A) site usage index; BP, biological process; KEGG: Kyoto Encyclopedia of Genes and Genomes.

Based on this division, the distal poly(A) site usage (PDUI) of each informative APA event between the two groups is compared. There are 2,779 transcripts displaying significantly lower PDUI values in mesenchymal phenotype, more than the transcripts presenting higher PDUI values for the NSCLC patients from TCGA (Figure 1D). As for the mesenchymal cell lines, the number of transcripts which show reduced PDUI values is 1,625. It is bigger than the number of transcripts that tell an opposite scenario (Figure 1D). Further, the average PDUI distribution of these differential APAs also reveals that the mesenchymal samples have significantly smaller PDUI values (Figure 1E). All these analyses describe that most transcripts prefer a shortened 3’-UTR after the transition from epithelial to mesenchymal status.

These differential APA events involve 3,350 genes for TCGA samples and 2,923 genes for CCLE samples. They are both enriched in EMT-related pathways (*P* < 0.05, FDR < 0.2, Figure 1F). For example, sphingolipids modulate EMT in cancer [16]. The tight junction is one connection way between epithelial cells and each other [4]. The neurotrophin receptor TrkB promotes lung adenocarcinoma metastasis [17]. These pathways describe the potential roles of APA events in the EMT process of NSCLC.

### 2. APA events have good prediction ability of EMT status for NSCLC

The 3,483 differential APA events in TCGA data (Figure 1D) are then screened using the pipeline shown in Figure 2A to identify 33 APA events highly associated with EMT. These APA events locate in lung cancer genes (Table S1) and may involve in the regulation of genes that are abnormally expressed after the transition from epithelial to mesenchymal status.

**Figure 2.**
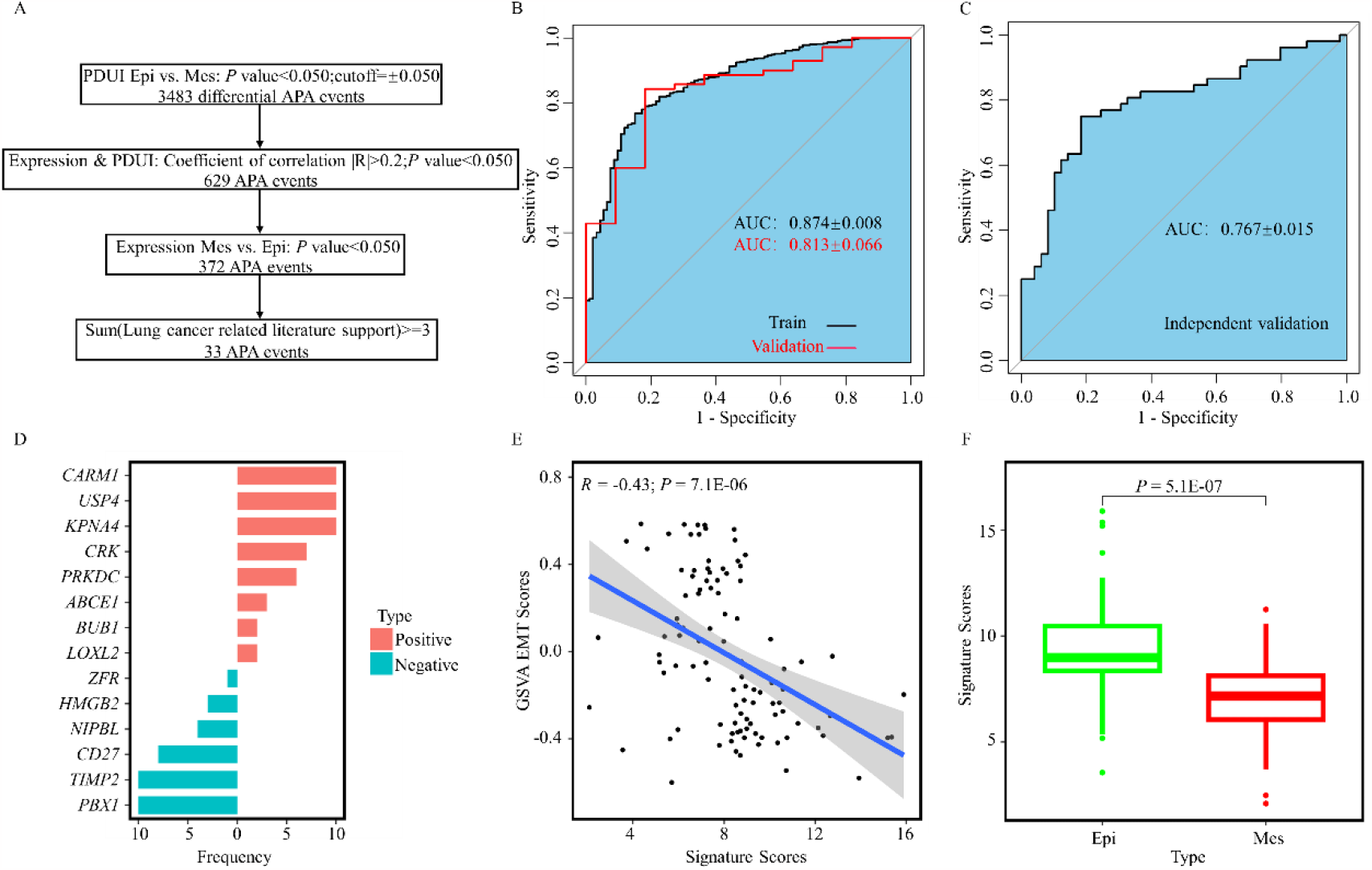
The identification of APA events related to EMT. (A) The pipeline to identify APA events related to EMT. (B) The ROC curve of logistic regression analysis on the 33 APA events for training and validation datasets of TCGA using 10-fold cross-validation method. (C) The ROC curve of logistic regression analysis on the 33 APA events for the independent CCLE dataset. (D) The number of the 14 APA events selected by the logistic regression model for prediction during the 10-fold cross-validation process. (E) The signature built on these 14 APA events is significantly associated with the score of EMT gene set by GSVA method for CCLE cell lines. The association for TCGA patients is shown in Figure S1B. (F) The significantly differential values of the APA signature between the defined epithelial and mesenchymal cell lines. The differential analysis for TCGA patients is shown in Figure S1C. EMT: epithelial-mesenchymal transition; ROC, receiver operating characteristic curve; AUC, area under the curve.

On these EMT-related APA events, the logistic regression method can provide a relatively accurate prediction of epithelial and mesenchymal status tested by 10-fold cross-validation (Figure S1A). The areas under the receiver operating characteristic curve of this model are 0.874 ± 0.008 for the training cohort of TCGA samples, 0.813 ± 0.066 for the validation cohort of TCGA samples (Figure 2B), and 0.767 ± 0.015 for the independent CCLE validation cohort (Figure 2C).

During the training process, 14 APA biomarkers participate in the prediction task (Figure 2D). A signature established with these 14 APA events for TCGA data shows significant associations with the score calculated on the EMT gene set from the Molecular Signatures Database (MSigDB) by the GSVA method (Figure S1B). Moreover, it has significantly differential APA score for the defined mesenchymal patients compared to that of epithelial patients (Figure S1C). The same 14 APA events and the coefficients are also used in cell line data to reveal the robustness of these EMT biomarkers (Figure 2E-F).

### 3. The proximal poly(A) selection of *CARM1* is a pathological biomarker of mesenchymal tumor

Of the 14 APA biomarkers, five APA events are highly relevant to EMT in all the ten times of cross-validation tests (Figure 2D). They are all associated with the expressions of corresponding APA genes. Specifically, only one gene, coactivator associated arginine methyltransferase 1 (*CARM1*), is dysregulated in all cancer patients and cell lines with mesenchymal status, tumors with higher clinical stages, and samples with poorer survival risks. It also plays an important role in the activation of TGF-β/ Smad pathway and the induction of EMT [18]. Thus, the APA event of this gene is selected to reveal APA-related mechanisms during the EMT process.

For this gene, the mesenchymal samples frequently select the proximal poly(A) site. This selection leads to the binding loss of miRNAs (Figure 3A), such as the EMT-related *miR-128* [19]. The lost miRNA regulation then causes the over-expressions of this gene (Figure 3B). One previous study describes the induction roles of *CARM1* in the EMT process [18]. Here, the higher expression of this gene is also discovered in more severe patients and cell lines characterized by EMT feature, clinical stage, and survival risk (Figure 3C). Thus, the proximal selection of poly(A) site of *CARM1* is suggested as a pathogenetic biomarker of NSCLC metastasis through the upregulation of *CARM1*. This hypothesis is also supported by the lower PDUI values in patients and cell lines with mesenchymal status, higher tumor stages, and poorer survival risks (Figure 3D).

**Figure 3.**
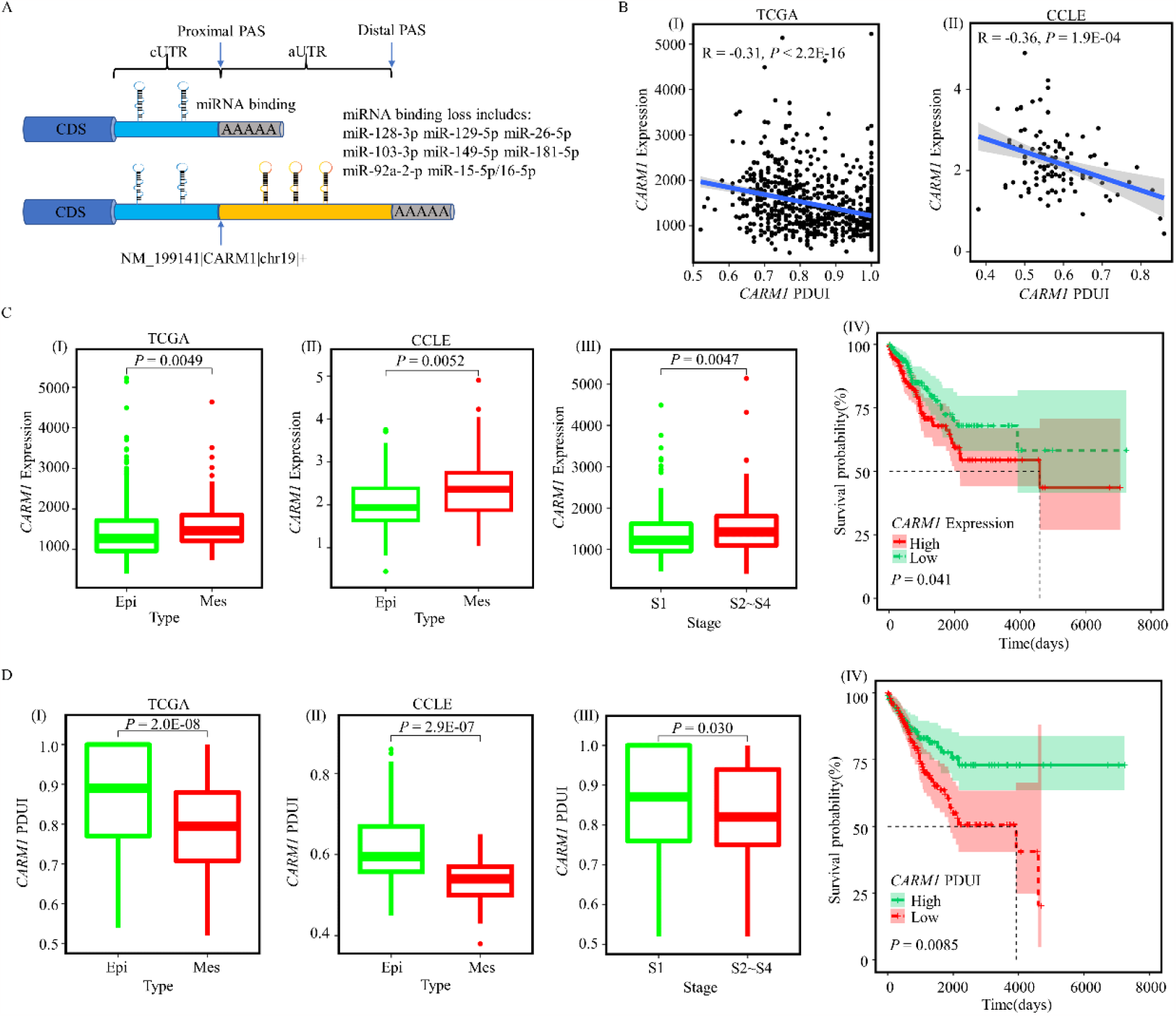
One potential mechanism for the associations between the APA event of *CARM1* and its own gene. (A) The diagram showing the proximal poly(A) selection of this APA event leads to the binding loss of miRNAs. (B) The associations between the PDUI values of the APA event and the expressions of *CARM1* for TCGA patients (left) and CCLE cell lines (right). (C) The comparison of *CARM1* expressions between epithelial and mesenchymal patients (left boxplot), between epithelial and mesenchymal cell lines (middle boxplot), between low and high clinical stages (right boxplot), and between low and high survival risks (right). (D) The comparison of the APA event of *CARM1* between epithelial and mesenchymal patients (left boxplot), between epithelial and mesenchymal cell lines (middle boxplot), between low and high clinical stages (right boxplot), and between low and high survival risks (right).

Further, the proximal poly(A) selection of *CARM1* can release miRNAs to regulate other genes. These genes have negative associations with *CARM1* expressions and positive correlations with the APA event of *CARM1* (Figure 4A). Of them, there are 279 genes involved in the biological processes related to EMT (Figure 4B). They include one RNA-binding protein, *RBM47*, which is reported as a suppressor of cancer progression and metastasis [20-21]. In lung cancer patients from TCGA and cell lines from CCLE, the reduced expression of *RBM47* is also discovered in samples with mesenchymal status, higher tumor stages, and poorer survival risks (Figure 4C).

**Figure 4.**
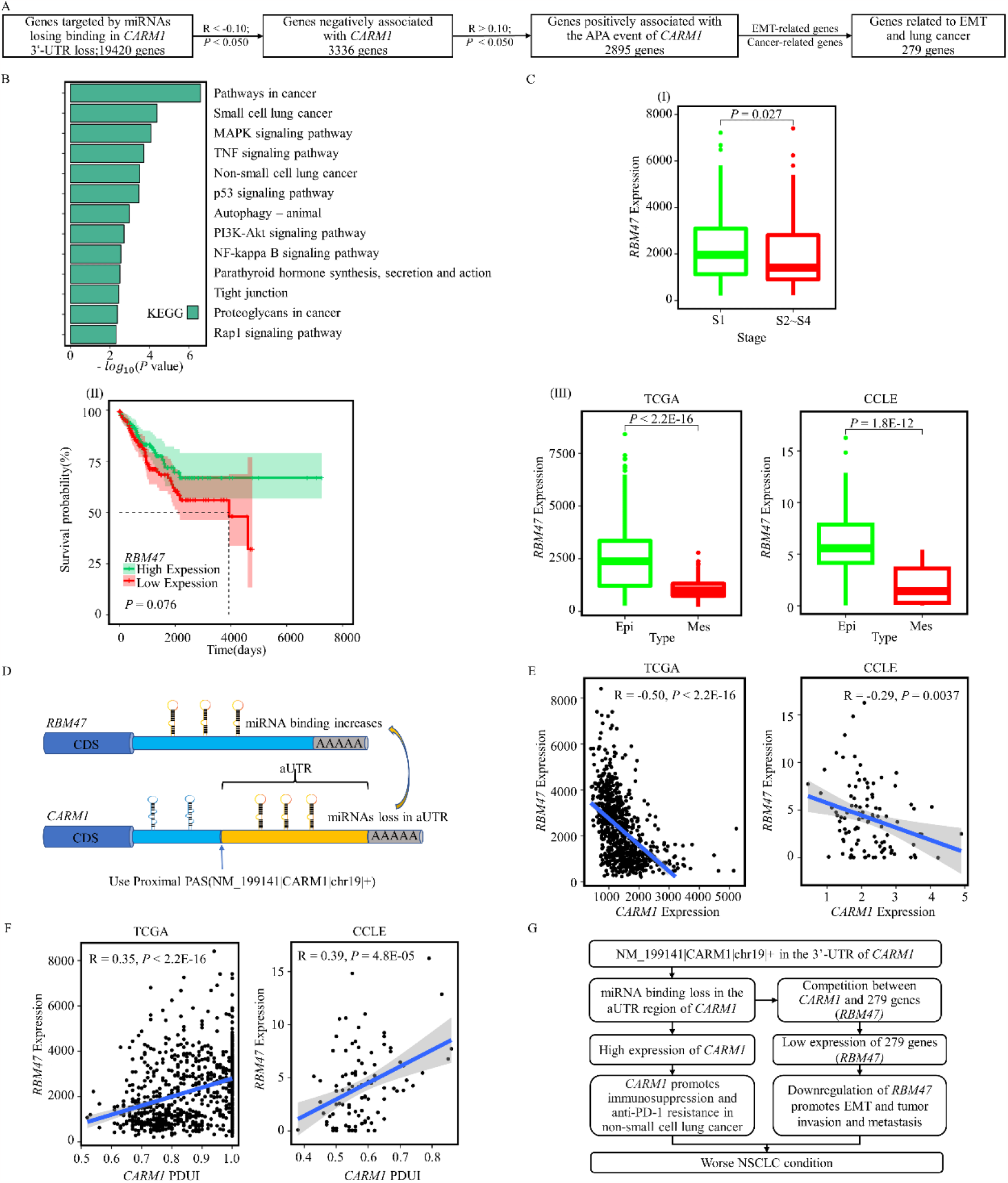
The APA event of *CARM1* may also regulate other EMT-related genes though interfering in miRNA regulations. (A) The pipeline for the identification of other genes related to EMT and lung cancer possibly affected by this APA event. (B) The biological functions possibly affected by this APA event. (C) The comparison of *RBM47* expressions between low and high clinical stages (I), between low and high survival risks (II), between epithelial and mesenchymal patients, and between epithelial and mesenchymal cell lines (III). (D) The diagram shows the altered miRNA regulations on *CARM1* and *RBM47* related to this APA event. (E) The expression associations between *CARM1* and *RBM47* for TCGA patients (left) and CCLE cell lines (right). (F) The associations between the PDUI values of this APA event and the expressions of *RBM47* for TCGA patients (left) and CCLE cell lines (right). (G) The hypothesis for the roles of this APA event in epithelia-mesenchymal transition and lung cancer progression.

The dysregulation of *RBM47* provides additional evidence about the pathology of the proximal poly(A) selection of *CARM1*. This selection releases the miRNAs to downregulate *RBM47* (Figure 4D). It is supported by the significantly negative expression correlations between *CARM1* and *RBM47* (Figure 4E) and the positive associations between *RBM47* and the APA event in both TCGA patients and cancer cell lines (Figure 4F). Thus, the proximal poly(A) selection of *CARM1* is a pathological biomarker of mesenchymal tumor and cancer metastasis through the upregulation of the EMT inducer of *CARM1* and the downregulation of the EMT inhibitor of *RBM47* (Figure 4G).

### 4. The proximal poly(A) selection of *CARM1* is associated with cisplatin drug responses

The above analyses reveal the involvement of the APA event of *CARM1* in the EMT process. Previous increasing evidence shows that EMT greatly impacts the responses of tumors to a broad number of anti-cancer drugs [22-23]. Thus, the effect of the APA event on drug sensitivity is studied.

There are two drugs interacting with *CARM1* collected from the CTD database [24]. They present lower IC50 (half maximal inhibitory concentration) values in mesenchymal samples (Figure 5A, Figure S2). It means that a low concentration of drugs is needed to inhibit tumor cells, representing higher drug sensitivities for mesenchymal tumors. The trend is consistent with the previous study showing that mesenchymal cells demonstrate greater sensitivity to certain chemotherapies [25]. These drugs include cisplatin, a chemotherapy medication for multiple tumors covering lung cancer.

**Figure 5.**
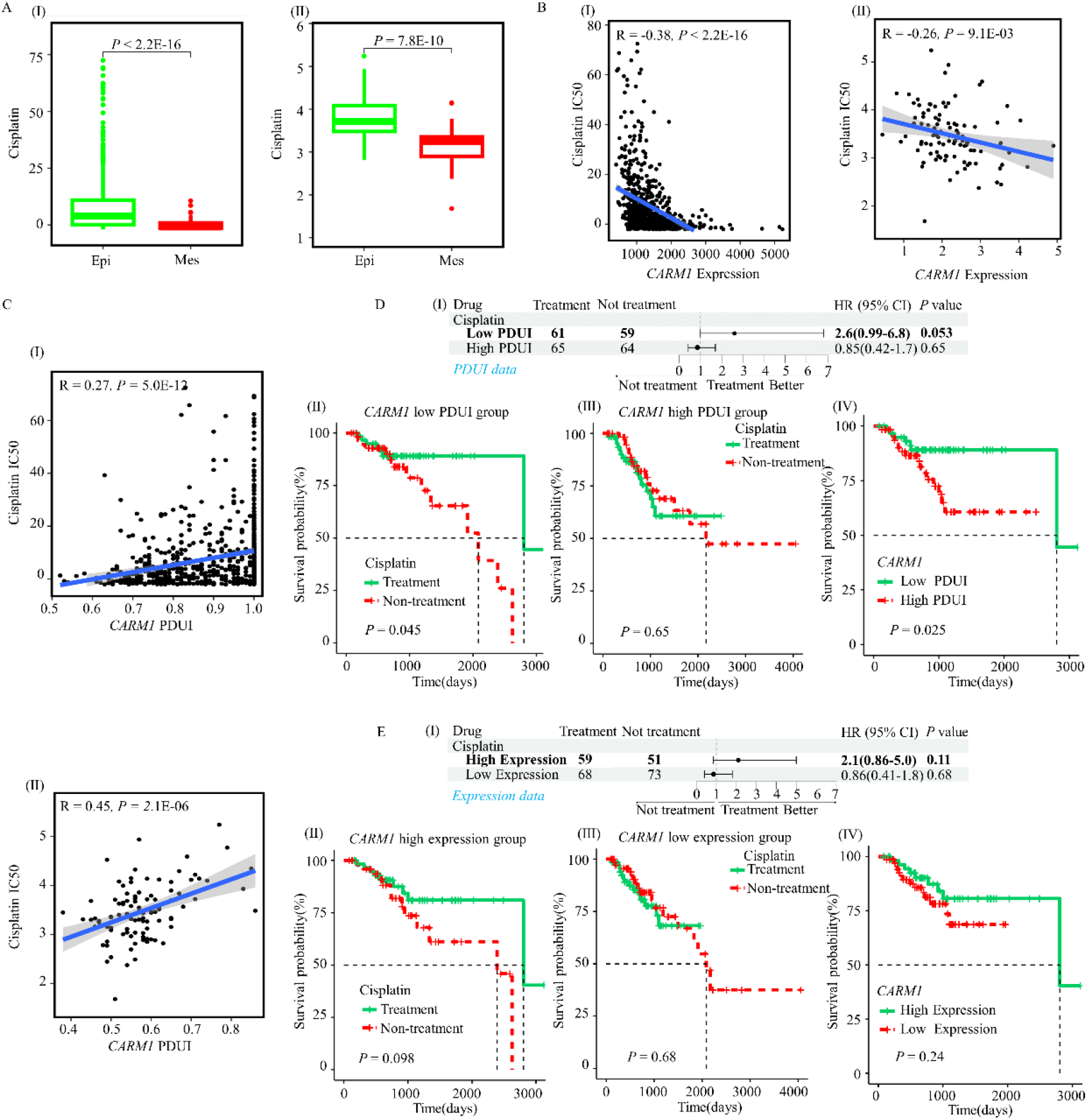
The associations of the APA event of *CARM1* with drug sensitivity. (A) The comparison of cisplatin sensitivity between the defined epithelial and mesenchymal patients (left) and cell lines (right). (B) The associations of cisplatin sensitivity with *CARM1* expression in TCGA patients (left) and CCLE cell lines (right). (C) The associations of cisplatin sensitivity with the APA event of *CARM1* in TCGA patients (top) and CCLE cell lines (bottom). (D) The Cox (top) and KM (bottom left) survival analysis between patients treated with cisplatin and patients untreated with cisplatin for two PDUI groups. One group includes patients with low PDUI value of the APA event of *CARM1*, while the other group contains patients with high PDUI value of the APA event of *CARM1*. The bottom right panel shows the KM survival analysis between low and high PDUI values for patients all treated with cisplatin. (E) The Cox (top) and KM (bottom left) survival analysis between patients treated with cisplatin and patients untreated with cisplatin for two expression groups. One group includes patients with high *CARM1* expression, while the other group contains patients with low *CARM1* expression. The bottom right panel shows the KM survival analysis between low and high *CARM1* expression for patients all treated with cisplatin. IC50, half maximal inhibitory concentration.

The IC50 of this drug and the expression of *CARM1* present negative associations in both patients and cell lines (Figure 5B). Higher expression of *CARM1*, lower values of IC50 and more sensitivity of tumors to this drug. Due to the involvement of the APA event in the miRNA regulations of *CARM1*, the proximal poly(A) selection of *CARM1* is associated with the higher cisplatin sensitivity (Figure 5C). Less concentration of cisplatin is needed to remove tumor cells for patients with the shortened 3’-UTR of *CARM1*.

The real and sufficient information for patients treated with cisplatin from TCGA also validates the associations of this APA event with drug sensitivity. This drug presents different effectiveness for patients with low and high PDUI values (Figure 5D). It seems that cisplatin can only improve the survival of patients with shortened 3’-UTR of *CARM1*. This hypothesis is also supported by the survival analysis for patients treated with cisplatin between proximal and distal ploy(A) selection groups. Specifically, this treatment design based on the poly(A) site is better than that with the expression of *CARM1* (Figure 5E). All the results reveal that the usage of cisplatin in clinics is required for personalized design base on the APA event of *CARM1*.

## Discussion

Our study analyzes the potential roles and underlying mechanisms of APA during the transition from epithelial to mesenchymal status in the pathogenesis and treatment of lung cancer. It expands the knowledge related to EMT and lung cancer from transcription factor [26-27], miRNA [28-29], protein [9, 30], and so on to APA. It supplements the crucial APA functions in the regulation of tumor genes [13, 15]. It also highlights the importance of APA for the treatment design of lung cancer patients [31-32].

During the study, at least two kinds of independent results are used to enhance the reliability of the analyses. For example, the definition of epithelial and mesenchymal groups is demonstrated with consistent features using different methods. The poly(A) site selection trend in mesenchymal samples, EMT-related APA biomarkers identification, and APA effect on drug responses are all validated in both datasets of lung cancer patients and cell lines. The estimated drug effectiveness is also verified in the patients with real treatment information. All these procedures aim to provide a rigorous and reliable conclusion.

Moreover, the shortened 3’-UTRs are also observed in mesenchymal cells (Figure S3A) after the analysis of single-cell RNA-seq data [33]. For this data, the epithelial and mesenchymal cells are annotated with the markers provided in that study and then inferCNV [34] (Figure S3B). This annotation is also verified with the score of the EMT gene set from MSigDB [35] (Figure S3C) and the pseudo-time trajectory result (Figure S3D). This analysis of single-cell RNA-seq provides additional evidence of the proximal poly(A) selection trend after the transition from epithelial to mesenchymal status.

The whole analysis points out one biomarker of NSCLC, the aberrant poly(A) selection of *CARM1*. It may affect tumor invasion, survival risk, and drug response by interfering in miRNA regulation on this tumor gene. Besides miRNAs, RNA-binding proteins (RBPs) are potentially involved in this process (Figure S4A-B). The selection of the proximal poly(A) site may also lead to the binding and regulation loss of RBPs, such as *ELAVL1* (Figure S4C-E). A further comprehensive analysis of RBP regulations interfered with APA is one future project.

## Methods

### 1. Sample preparation

There were 1,013 NSCLC patients from TCGA and 123 NSCLC cell lines from CCLE [36] involved in this study for APA effects on the transition from epithelial to mesenchymal status. The corresponding clinical information of these samples included relapse-free survival (RFS), tumor stage, treatment drugs, and so on.

### 2. Detection of APA events

The APA events of patients from TCGA were directly downloaded from The Cancer 3’UTR Atlas (TC3A) [37]. As for the APA events of cell lines, a similar procedure as TC3A was used. Specifically, samtools (Version: 1.9) extracted sequences and qualities from RNA-seq bam files. STAR (Version: 2.7.9a) [38] aligned the reads to the hg19 reference genome (Gencode v37lift37). DaPars [39] was applied to identify APA events and quantify PDUI values. Informative APA events were recognized as events with missing values of less than 60% and consistent one or zero PDUI values of less than 40%. Eventually, there were 6,914 informative APA events of patients and 16,259 informative APA events of cell lines retained in the following analyses.

### 3. Quantification of gene expression

The gene expression of patients was directly downloaded from TCGA (hg19). As for the gene expression (Upper Quartile Normalization, UQN) of cell lines, the similar reference genome and procedures as TCGA were used with RSEM [40]. The low abundant genes with mean expressions of less than one UQN were not considered in the following analysis, reserving 17,735 informative genes for patients and 7,899 informative genes for cell lines.

### 4. Definition of epithelial and mesenchymal status

For each NSCLC patient from TCGA, a two-sample Kolmogorov-Smirnov test on 315 genes was used to assess the status of epithelial or mesenchymal phenotype [41]. For a cell line, the same method on 218 genes from the same literature was applied. A sample with a positive score (*P* < 0.05) was defined as mesenchymal NSCLC, while a sample with a negative score (*P* < 0.05) was recognized as epithelial NSCLC. The division of all samples was visualized by principal component analysis (PCA) [42].

Then the enrichment scores of three EMT-related gene sets using the GSVA [43] method were compared between the defined epithelial and mesenchymal groups to verify the reliability of the definition. These three gene sets were EPITHELIAL_MESENCHYMAL_TRANSITION, HYPOXIA, and TGF_BETA_ SIGNALING from The Molecular Signatures Database (MSigDB) [35]. The hypoxia microenvironment played a role in the induction of EMT-related genes [44]. The TGF-β signaling pathway participated in the metabolism processes of glycolysis and mitochondrial respiration, whose consequences for the redox state and lipid metabolism were in favor of EMT [45].

### 5. Identification of APA biomarkers related to EMT

The EMT-related APA biomarkers were defined as events showing differential usage of poly(A) sites in mesenchymal tumors (*P* < 0.05 and |log2FC| > 0.05) and significant associations with EMT-related genes (*P* < 0.05 and |R| > 0.2).

The EMT-related genes were recognized by their abnormal expressions in the mesenchymal group (*P* < 0.05) and their functions related to lung cancer from previous studies (Table S1). At last, a logistic regression model was used to select the most significant APA biomarkers associated with EMT. Their effectiveness was assessed by the area under the receiver operating characteristic curve, correlations with the EMT enrichment scores, and differences between the defined epithelial and mesenchymal groups.

### 6. Analysis of APA-mediated miRNA regulation during EMT process

The selection of poly(A) sites can alter the lengths of 3’-UTRs and disturb miRNA regulations on genes. According to TargetScan (v7.2) [46], the proximal poly(A) usage may cause the loss of conserved miRNA binding targets. The lost miRNA regulations were verified with the negative associations between PDUI values of the APA event and expressions of the APA genes. Further, this proximal poly(A) selection may not only upregulate the APA gene but also release miRNAs to dysregulate other genes. The APA effect on other genes was supported by their positive associations combined with the negative associations of the two genes.

### 7. Analysis of APA effect on drug sensitivity

The APA events affected the migration ability of tumor cells, which may alter the effectiveness of drug treatment. This study started with the identification of potential drugs targeting the genes with the APA biomarkers from the CTD database [24]. The sensitivity of these drugs on cancer patients was evaluated by pRRophetic [47] with IC50 index. The positive association between PDUI values and drug sensitivity indicated that patients with shortened 3’-UTR of the APA event would have better drug responses. It was then verified in TCGA patients with real treatment information through the comparison of survival risks and drug treatment effectiveness between low and high PDUI cohorts.

## Acknowledgement

The results <published or shown> here are in whole or part based upon data generated by the TCGA Research Network: https://www.cancer.gov/tcga. This research was funded by the National Natural Science Foundation of China (Grant No. 62002270), the Fundamental Research Funds for the Central Universities, the National Natural Science Foundation of China (Grant No. 82227802), the Natural Science Foundation of Shaanxi Province of China (Grant No. 2020JQ-332), the China Postdoctoral Science Foundation (Grant No. 2018M643583), and the National Key R&D Program of China (Grant No. 2017YFA0205202) and partially funded by the National Natural Science Foundation of China (Grant No. 61672422).

## Informed Consent Statement

The data was obtained from public resources including The Cancer Genome Atlas (TCGA) and Cancer Cell Line Encyclopedia (CCLE).

## Data Availability Statement

The data presented in this study are available in Supplementary Material here.

## Author Contributions

Conceptualization, S.W.; methodology, S.W. and X.Q.; formal analysis, X.Q.; investigation, S.W. and X.Q.; data curation, X.Q; writing—original draft preparation, S.W., and X.Q.; writing—review and editing, L.H.; supervision, L.H.; funding acquisition, S.W. and L.H. All authors have read and agreed to the published version of the manuscript.

## Conflicts of Interest

The authors declare no conflict of interest. The funders had no role in the design of the study; in the collection, analyses, or interpretation of data; in the writing of the manuscript; or in the decision to publish the results.

## Notes

### Competing Interest Statement

The authors have declared no competing interest.

