## Supplementary Table 1 and Figure 1-4 for "The role of alternative polyadenylation in Epithelial-mesenchymal transition of non-small cell lung cancer"

TableS1 The genes related to lung cancer according to previous studies

| Gene | Pubmed ID |
|---|---|
| *MAP3K2* | 35929837,34484674,27498924 |
| *PARP1* | 37066722,37060095,36919822 |
| *KPNA4* | 36573325,33407505,32151944 |
| *CRK* | 27027347,25956913,24167658 |
| *MYH9* | 36268014,34761592,34635641 |
| *TOPBP1* | 28628491,27729767,26003539 |
| *ATF2* | 35692887,33008584,32476275 |
| *USP4* | 36762777,35027535,32549341 |
| *CD68* | 37035471,35928634,33392661 |
| *PML* | 36476834,30532072,28744813 |
| *EEF2* | 32943920,32054489,30147338 |
| *DDR1* | 32759499,31979355,31659178 |
| *PRKDC* | 32238472,25823795,25495526 |
| *CARM1* | 36690626,35664694,34052308 |
| *ZFR* | 33407505,31407591,30611568 |
| *ROCK1* | 35984492,35626071,31986487 |
| *NIPBL* | 33660799,29670369,33660799 |
| *GPNMB* | 33706413,31822499,26883195 |
| *BUB1* | 36309619,35117845,11146226 |
| *P4HA1* | 34659558,34049151,33299876 |
| *TPX2* | 37009264,36389338,32748897 |
| *ABCE1* | 33537917,31841188,31306106 |
| *PIK3CA* | 36898331,36835617,36644157 |
| *LOXL2* | 35483499,29156517,27694892 |
| *HMGB2* | 36445573,36416408,34651416 |
| *MPG* | 28489575,25355292,16613673 |
| *PBX1* | 36361531,33113625,25400423 |
| *CD151* | 34108040,33428143,32117989 |
| *TRIM27* | 36200036,34932879,34695898 |
| *MEF2D* | 34290714,29963192,28498474 |
| *PPDPF* | 35906391,34975328,30954221 |
| *TIMP2* | 34732411,34178128,31632575 |
| *CD27* | 34570961,34503105,31533124 |

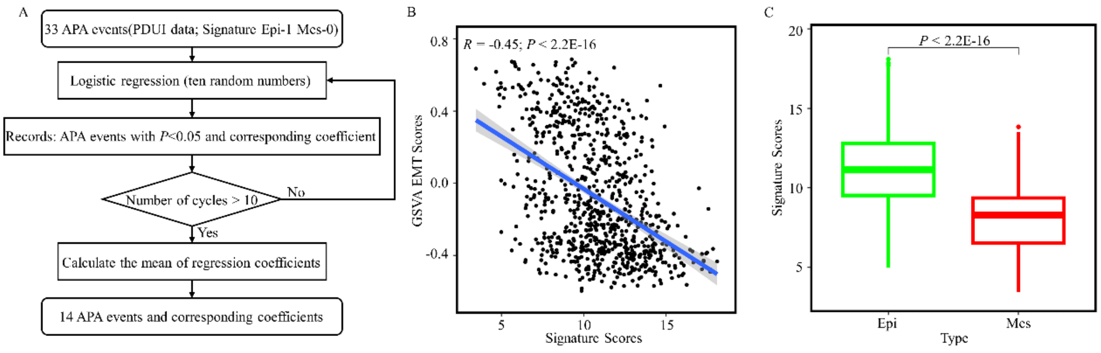


Figure S1. The supplemented identification process of APA events related to EMT. (A) The detail procedure of logistic regression method to select 14 APA events. (B) The signature built on these 14 APA events is significantly associated with the score of EMT gene set by GSVA method for TCGA patients. (C) The significantly differential values of the APA signature between the defined epithelial and mesenchymal patients.

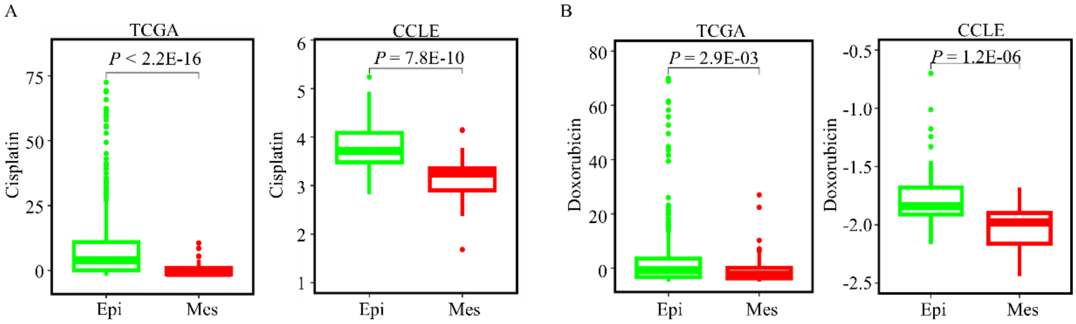


Figure S2. The EMT effect on drug sensitivity. (A) The comparison of cisplatin sensitivity between the defined epithelial and mesenchymal patients (left) and cell lines (right). (B) The comparison of doxorubicin sensitivity between the defined epithelial and mesenchymal patients (left) and cell lines (right).

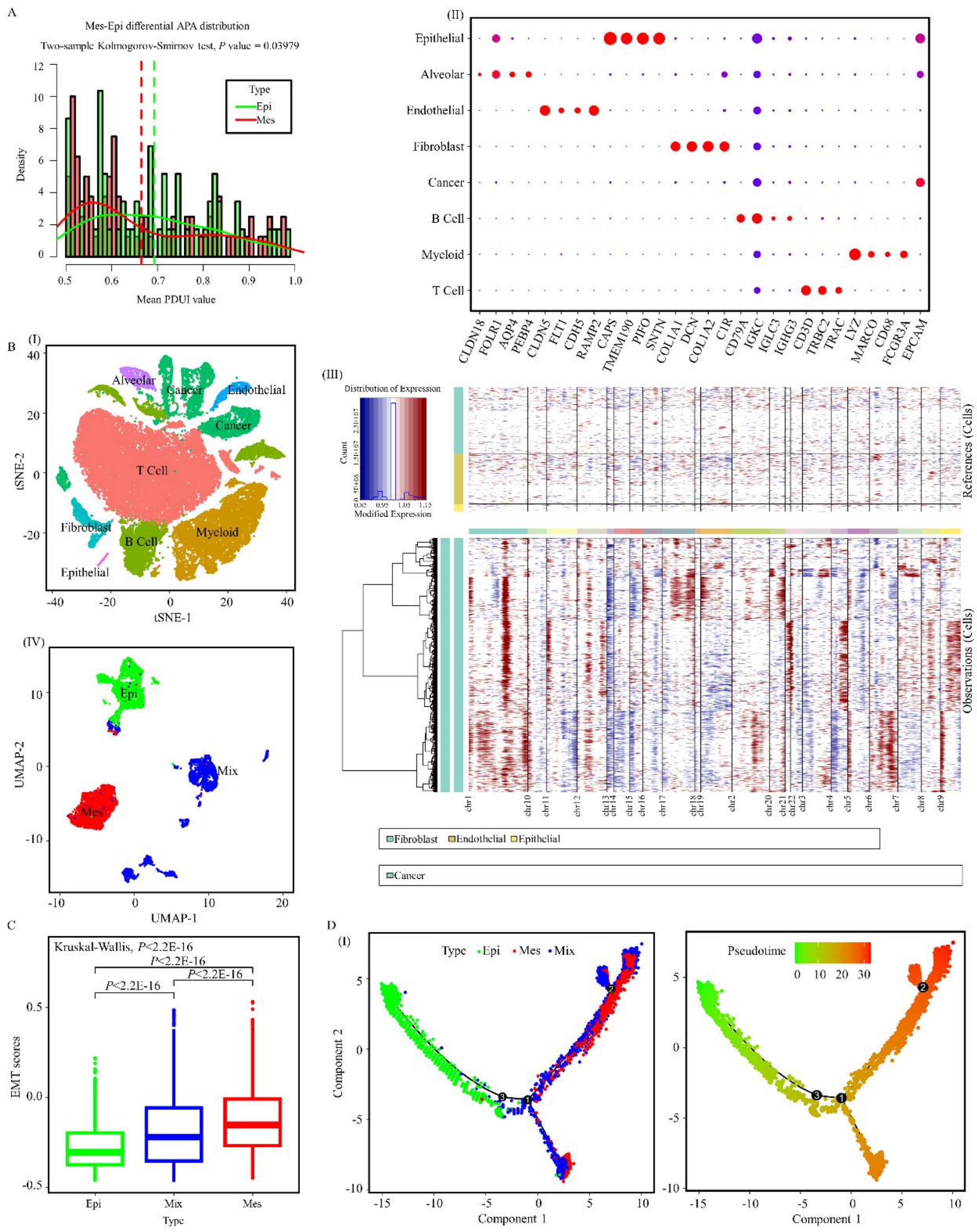


Figure S3. The analysis of alternative polyadenylation between epithelial and mesenchymal cells. (A) The PDUI distributions of average differential APA events in epithelial and mesenchymal cells. (B) The top left panel shows the clusters of the nine cell types. The top right panel shows the markers to annotate the nine cell types. The bottom left panel presents the annotation of epithelial and mesenchymal cells by inferCNV results shown in bottom right panel. (C) The comparison of the scores of EMT gene set between epithelial and mesenchymal cells. (D) The pseudo-time trajectory analysis of epithelial and mesenchymal cells.

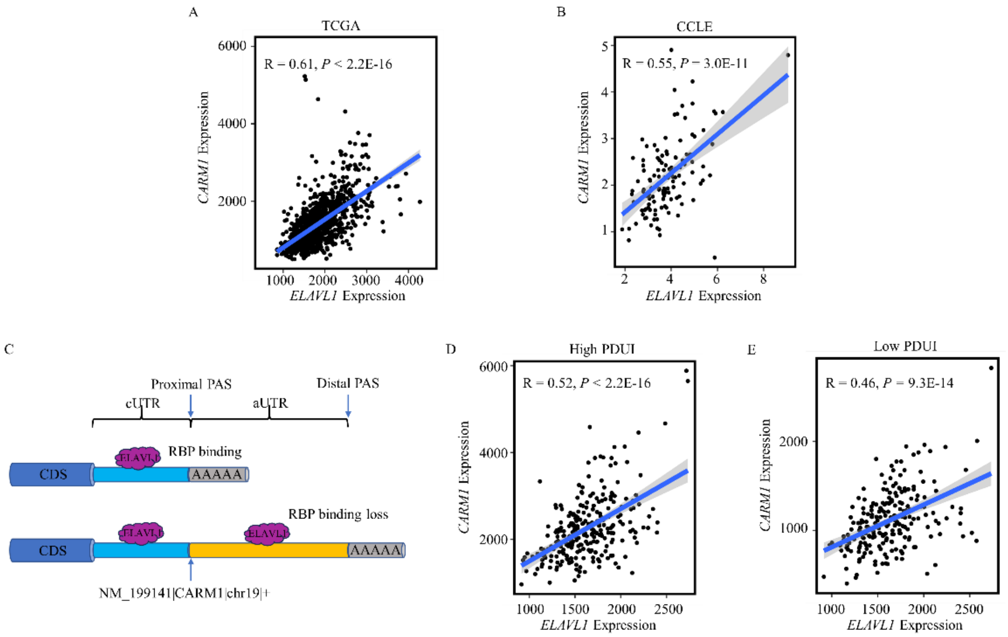


Figure S4. The effect of alternative polyadenylation on the RNA binding regulation of genes. (A) The associations between an RNA binding protein (*ELAVL1*) and *CARM1* gene expression in lung cancer patients. (B) The associations between an RNA binding protein and *CARM1* gene expression in lung cancer cell lines. (C). The proximal poly(A) selection leads to the binding loss of RNA binding proteins, including *ELAVL1*. (D) The associations between an RNA binding protein and *CARM1* gene expression in distal poly(A) selection group. (E) The associations between an RNA binding protein and *CARM1* gene expression in proximal poly(A) selection group.
